# Assembly of chromosome-scale contigs by efficiently resolving repetitive sequences with long reads

**DOI:** 10.1101/345983

**Authors:** Huilong Du, Chengzhi Liang

**Affiliations:** State Key Laboratory of Plant Genomics, Institute of Genetics and Developmental Biology, Chinese Academy of Sciences, 1 Beichen West Road No. 2, Beijing 100101, China; College of Life Sciences, University of Chinese Academy of Sciences, Beijing 100049, China

## Abstract

Due to the large number of repetitive sequences in complex eukaryotic genomes, fragmented and incompletely assembled genomes lose value as reference sequences, often due to short contigs that cannot be anchored or mispositioned onto chromosomes. Here we report a novel method **H**ighly **E**fficient **R**epeat **A**ssembly (HERA), which includes a new concept called a connection graph as well as algorithms for constructing the graph. HERA resolves repeats at high efficiency with single-molecule sequencing data, and enables the assembly of chromosome-scale contigs by further integrating genome maps and Hi-C data. We tested HERA with the genomes of rice R498, maize B73, human HX1 and Tartary buckwheat Pinku1. HERA can correctly assemble most of the tandemly repetitive sequences in rice using single-molecule sequencing data only. Using the same maize and human sequencing data published by Jiao *et al.* (2017) and Shi *et al.* (2016), respectively, we dramatically improved on the sequence contiguity compared with the published assemblies, increasing the contig N50 from 1.3 Mb to 61.2 Mb in maize B73 assembly and from 8.3 Mb to 54.4 Mb in human HX1 assembly with HERA. We provided a high-quality maize reference genome with 96.9% of the gaps filled (only 76 gaps left) and several incorrectly positioned sequences fixed compared with the B73 RefGen_v4 assembly. Comparisons between the HERA assembly of HX1 and the human GRCh38 reference genome showed that many gaps in GRCh38 could be filled, and that GRCh38 contained some potential errors that could be fixed. We assembled the Pinku1 genome into 12 scaffolds with a contig N50 size of 27.85 Mb. HERA serves as a new genome assembly/phasing method to generate high quality sequences for complex genomes and as a curation tool to improve the contiguity and completeness of existing reference genomes, including the correction of assembly errors in repetitive regions.

## Introduction

Assembly of high-quality genome sequences is an important task which is the basis for genome structure and evolutionary studies, and for gene mapping and cloning. Complex eukaryotic genomes contain a large number of repetitive sequences which complicates the genome assembly process^1^. In the past decade, next generation sequencing assisted the assembly of hundreds of draft animal and plant genomes. Improvements in sequencing read lengths to tens of kb by single molecule sequencing (SMS) technologies from Pacific Biosciences (PacBio)^2^ and most recently from Oxford Nanopore^3^ have enabled the assembly of many complex eukaryotic genomes. However, these assemblies are still fragmented and generate incomplete draft genomes usually consisting of thousands of contigs with many unresolved regions composed of long stretches of repetitive sequences^4–6^. The missing repeats in a genome assembly may have important functional implications. For example, segmental duplications account for ∼5% of the human genome and are associated with structural variations and genetic diseases^7^.

The SMS long reads are currently assembled using String Graph (SG)^8^ or -like assemblers such as FALCON^9^ and Celera Assembler^10^ (CA). CA represents the traditional Overlap-Layout-Consensus (OLC) assemblers and uses overlap graph^11^ to store reads and sequence overlaps between them. OLC is now used in several genome assemblers including PBcR^12^, CANU^13^ and MECAT^14^. A string graph^8^ reduces an overlap graph using a transitive reduction step to collapse identical or similar sequences. Tiling paths of reads are identified and joined together to form contigs, but multiple copies of similar repeats are also compressed. The approach assembles unique sequences reliably but repeats longer than the read length lead to branching paths and thus form fragmented contigs. Therefore, accurate assembly and resolution of large repeats and highly similar haplotypes in heterozygous genomes still remains a major challenge in genome assembly efforts.

Genome assembly projects often generate chromosome-scale pseudomolecules to anchor the assembled contigs/scaffolds with genetic maps or Hi-C data^15^. However, the approach often leaves many unordered and unoriented contigs, as well as unanchored orphan contigs. The genome maps from BioNano Genomics^16^ covering megabases (Mb) in length can be used to link contigs to hybrid scaffold sequences. However, the method cannot improve the contig lengths and often leaves unfilled gaps of hundreds of kb and many unmapped contigs due to lack of labelling enzyme recognition sites.

Therefore, our work is focused on resolving the repeats in genome assembly. With the current availability of long reads, repeats that are shorter than the read length, say 20 kb, do not usually cause assembly problems, and only the repeats that are longer than the read length need to be resolved. In our previous work^5^, we used fosmid clone sequences to assist the assembly of SMRT sequencing data. However, the relatively high cost of construction and sequencing of a fosmid library inhibits the wide application of this method to a large number of genomes. Here we report a highly efficient genome assembly method using SMS data to resolve repeats called HERA (**H**ighly **E**fficient **R**epeat **A**ssembly), which enables the assembly of highly contiguous genomes by assembling each individual repeat separately and correctly connecting it to its flanking sequences. BioNano genome maps and chromosomal anchoring information can help to resolve highly similar repeats. We tested the method on the genomes of rice, maize, human and Tartary buckwheat, and it dramatically improved on the sequence contiguity compared with the assemblies produced by existing assemblers. We generated a high-quality reference genome for maize and Tartary buckwheat with filled gaps and error corrections.

## Results

### Overview of HERA in assembling repeats

The algorithms used by HERA are fully described in the Methods. In this work, HERA constructs an overlap graph using the contigs generated by other assemblers such as PBcR, CANU, MECAT or FALCON as anchoring nodes and the corrected reads as extending nodes (Figure 1a, b). The contigs represent a non-redundant set of predefined sequences in a genome. Note that other type of sequences such as selected reads can also be used as anchoring nodes. By graph traversal under the same extending criteria, a set of paths (representing tiling read paths) are created between a starting anchoring node and a set of ending anchoring node (Figure 1c). By counting the number of paths between each pair of anchoring nodes, the overlap graph is reduced to a connection graph, where different repeat copies are used as separate connecting edges between anchoring nodes (Figure 1d). Among all the paths, HERA identifies the correct one and thus assembles the sequences between the true neighboring pairs of anchoring nodes into contigs (which also include the flanking sequences). The assembly of single-copy-only sequences between anchoring nodes is trivial. For repetitive sequences, HERA fully takes advantage of the sequence variations between different repeat copies to distinguish them from each other wherever possible (Figure 1a and Supplementary Figure 1). During the path extension process, the reads originating from the same repeat copy usually have a higher chance of being selected due to their higher sequence similarity in overlapping regions than those from different repeat copies. As a result, the sequences flanking a repeat will be connected by valid paths consisting of the reads originating from the same repeat and thus the correct sequences will be assembled.

**Figure 1:**
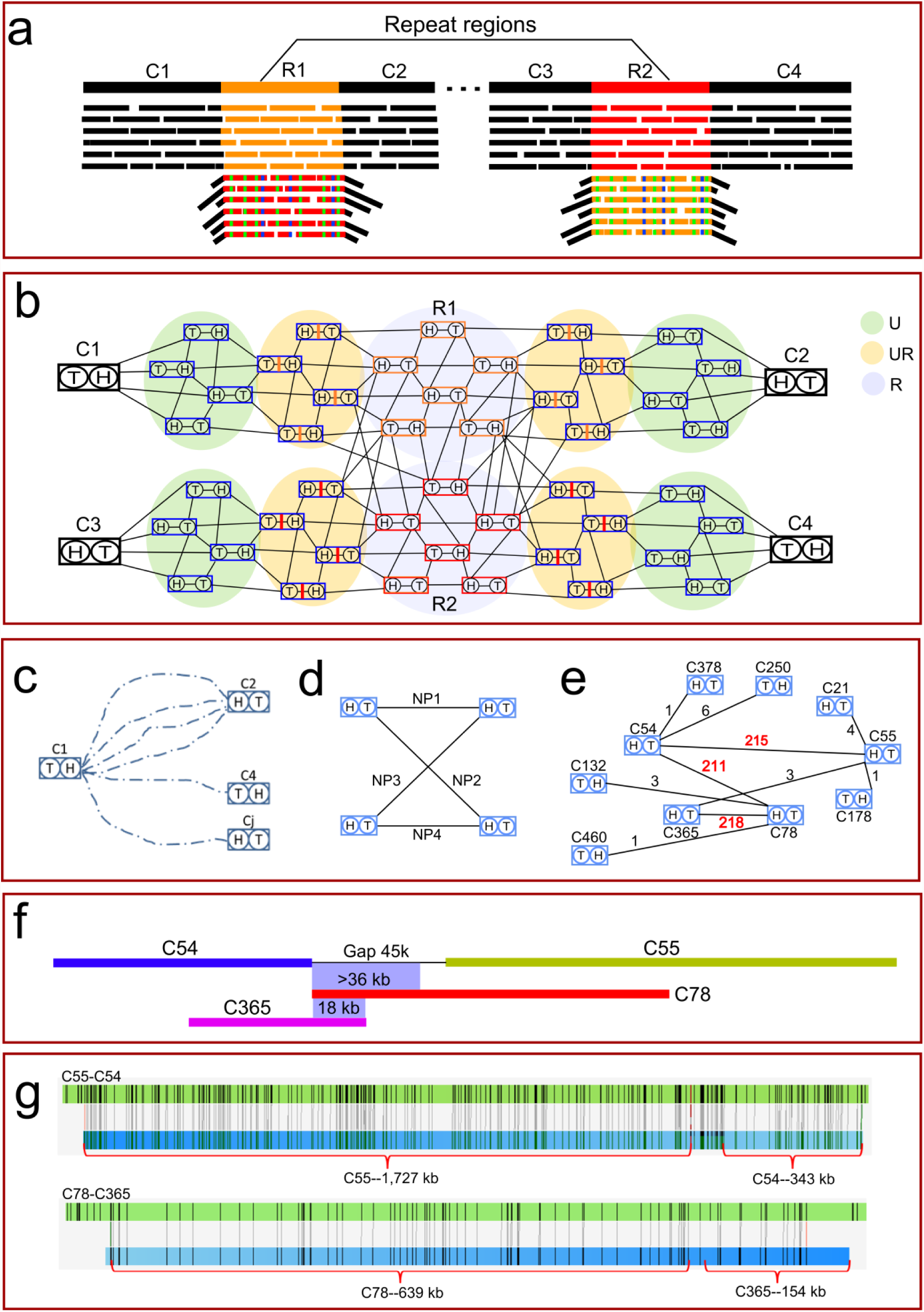
Overview of HERA. (**a**) Two copies of repeats (R1 and R2) are similar to each other but they also contain sequence variations which can be found in the reads originating from them. The alignments of junction reads (across the boundary between repeat and unique sequences) to a different repeat copy form overhangs of unaligned, unique sequences. (**b**) A subgraph of an overlap graph corresponding to the genome segments and sequencing reads shown in (a). The sequencing reads can be classified into three types: unique reads (U), repeat reads (R) and junction reads (UR). The edges are shown for illustration only; they may not be assigned correctly in this graph. (**c**) The path extension from contig end C_1_^h^ can reach a set of other contig ends, which include C_2_^h^, the true target, and C_4_^t^, the false target, and possibly others (C_j_^h^) from background noise. (**d**) A connection graph showing the number of paths (NP) between each pair of contig nodes. (**e**) A subgraph of a connection graph with examples of conflicting connections. The conflicting indices of two contig ends were: CI_54_^t^ = 211/215 = 0.98; CI_78_^h^ = 211/218 = 0.97. These conflicting connections can be resolved because the number of paths between C_365_^t^-C_55_^h^ was very small, so that C_78_^h^-C_365_^t^ can be connected first. (**f**) Sequence alignments showing a fragment of at least 36 kb in C_78_ being similar to the connecting sequence between C_55_^h.^ and C_78_^h^ and 18 kb of highly similar sequence in C_365_^t^ overlapping C_78_^h^. (**g**) The alignments to BioNano genome maps confirmed that the connections of C_54_^t^-C_55_^h^ and C_78_^h^-C_365_^t^ were correct.

Valid paths usually have the highest scores among all paths even after considering sequencing errors. For two nearly identical repeat copies, it is expected that the reads originating from them will be mixed with each other during the path extension process. In such a case, it is also expected that the number of valid paths to connect a pair of truly adjacent anchoring nodes and the number of invalid paths to connect a pair of non-adjacent anchoring nodes will be similar. This leads to conflicts in the connection graph (Figure 1e,f). There are several ways to resolve the conflicts: First, by removing one of the connected anchoring nodes when it can be more confidently connected to another anchoring node (Figure 1e); Second, by using BioNano genome maps or mate-pair scaffolding information to identify the correct pair (Figure 1g); Third, by using the chromosomal grouping information derived from genetic map or Hi-C data to identify the correct pair.

Due to both sequencing errors and sequence variations between the reads originating from different copies of repeats, not all connecting paths between the same pair of anchoring nodes have the same length. Further, the paths from tandem repeats or complex repeats can have very different lengths due to inclusion of different repeat units, which lead to multiple peaks in the length distribution plot (Figure 2). The difference between adjacent peaks corresponds to a repeat unit with possibly additional sequences. The distribution of the path lengths can be used to estimate the length of the repeat and the path sequences can be used to determine whether the repeat contains sequences other than the tandem units.

**Figure 2:**
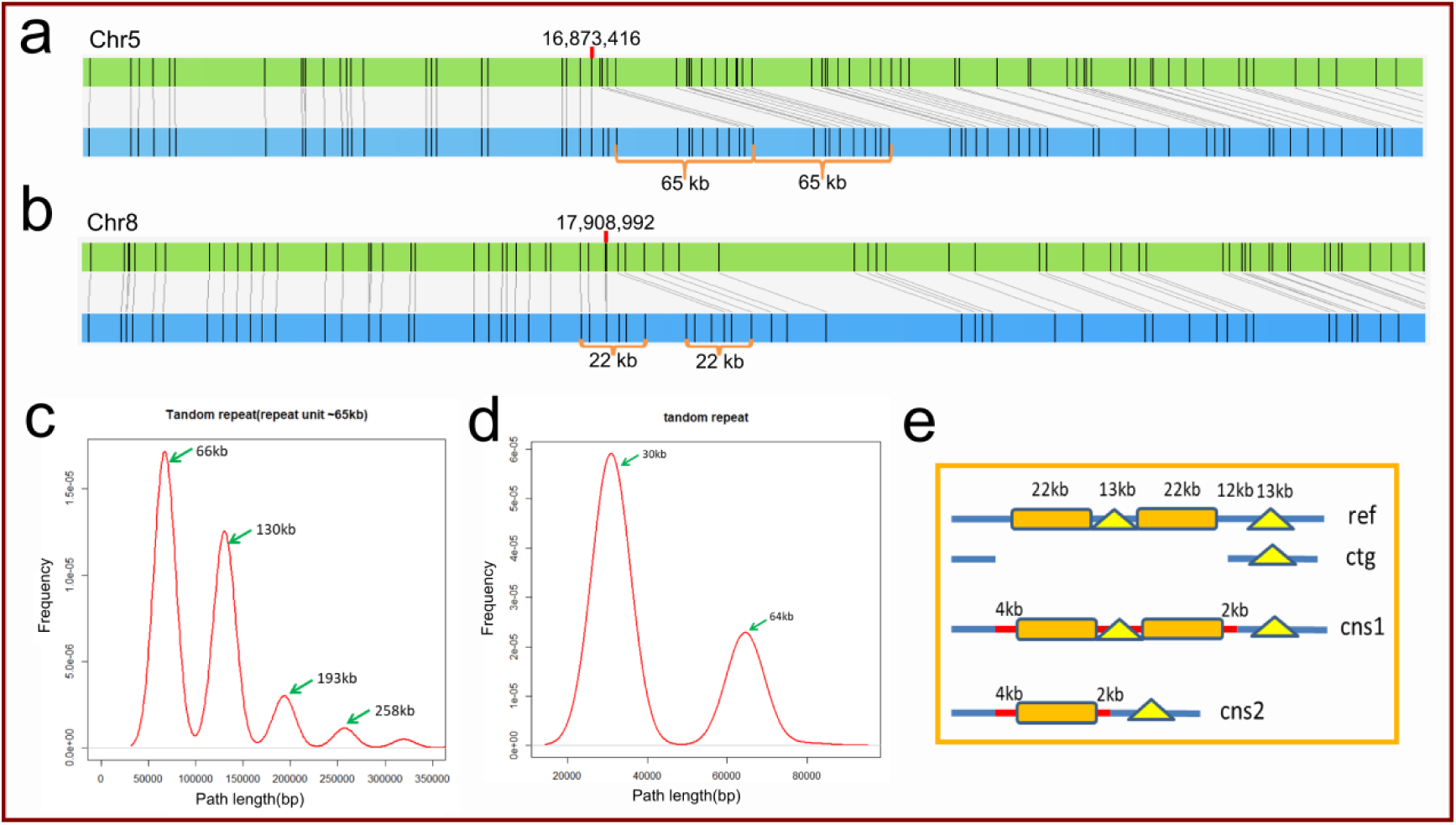
Assembly of tandemly repetitive sequences by HERA. (**a**) A tandemly repetitive sequence on chromosome 5 of R498 with a unit length of 65 kb. The upper green horizontal bar represents the genome sequence and the lower blue bar represents the BioNano map. (**b**) A repetitive sequence on chromosome 8 of R498 with a unit length of 22 kb. (**c**) The length distribution of HERA generated tiling paths for the repeat shown in (a). The paths are divided into several clusters and the distances between adjacent peaks are 65 kb which matched the repeat unit length in (a). The second peak represents the whole region of two repeat units (130 kb). (**d**) The length distribution of paths between flanking contigs in (b). The paths are divided into two clusters and the distance between the two peaks is around 35 kb. (**e**) The schematic representation of the repeat in (b). In this region, there are two repeat units being separated by a dissimilar repeat unit of 13 kb. Ref, the repeat including two units of 22 kb sequences; ctg, the flanking contigs to be connected; cns1 and cns2, excluding the two contig sequences shown in ctg, represent the second and the first peak in (d), respectively.

HERA is also extremely useful to fill gaps in hybrid genome maps or scaffolds, or fill gaps and fix errors in the pseudomolecules derived from genetic maps or Hi-C data. The known gap length is helpful for selecting the correct path. The connecting information (number of paths and overlap sequence similarity) between contigs is helpful to determine the order and orientation of the contigs in pseudomolecules.

### Validation of HERA based on assembly of the rice R498 genome

We tested HERA (Supplementary Figure 2) using the SMRT sequencing data of R498^5^ (Supplementary Table 1). First, we used CANU to assemble the genome into 402.5 Mb sequences with a contig N50 size of 1.3 Mb (Figure 3 and Supplementary Table 2). After HERA assembly, the contig N50 size was increased to 13.2 Mb. With the help of BioNano genome maps, which resolved a total of 61 conflicting connections, the contig N50 size was further increased to 14.4 Mb, with chromosome 8 assembled to a single contig. After non-rice genome sequences were filtered out by comparing them to the R498 reference genome^5^ (R498_ref1), only 40 contigs were left in the HERA assembly to form pseudomolecules, reaching a total genome size of 391.6 Mb. For simplicity, we did not try further to fill the gaps left in the pseudomolecules. The HERA assembly was approximately 1.5 Mb larger than R498_ref1. Comparison between the assemblies showed that 95% of the newly increased sequences were centromeric or subtelomeric repeats. Notably, the CANU assembly contained several chimeric contigs which were fixed since they were easy to identify in the HERA assembly due to the increased contig length, by comparison to genome maps or by using the genetic map we constructed previously^5^.

**Figure 3:**
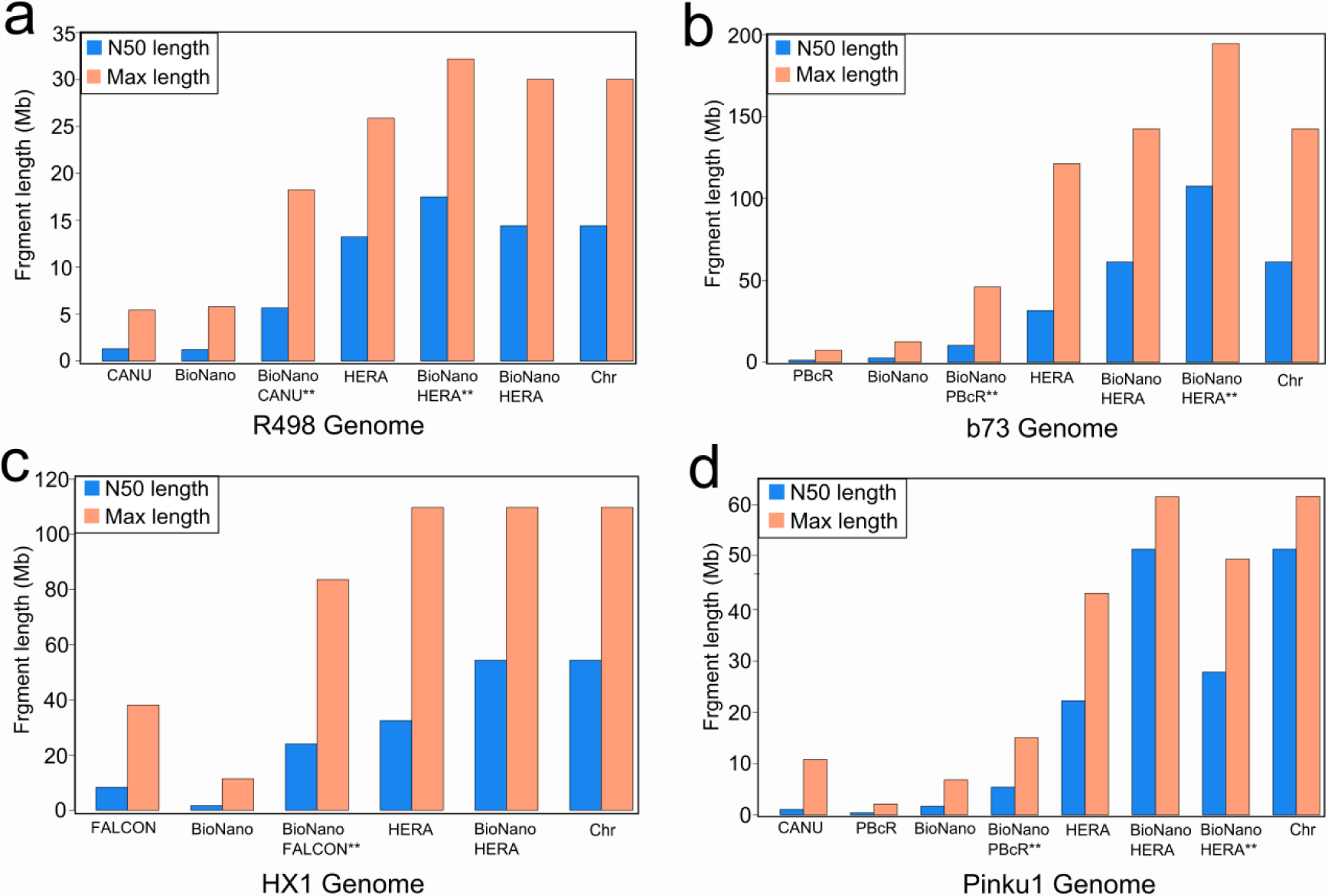
Statistics of genome assemblies of rice, maize, human and Tartary buckwheat. BioNano, genome maps. **indicates hybrid scaffolds. All other sequences are contigs. Chr, the HERA contigs anchored to chromosomes.

**Figure 4:**
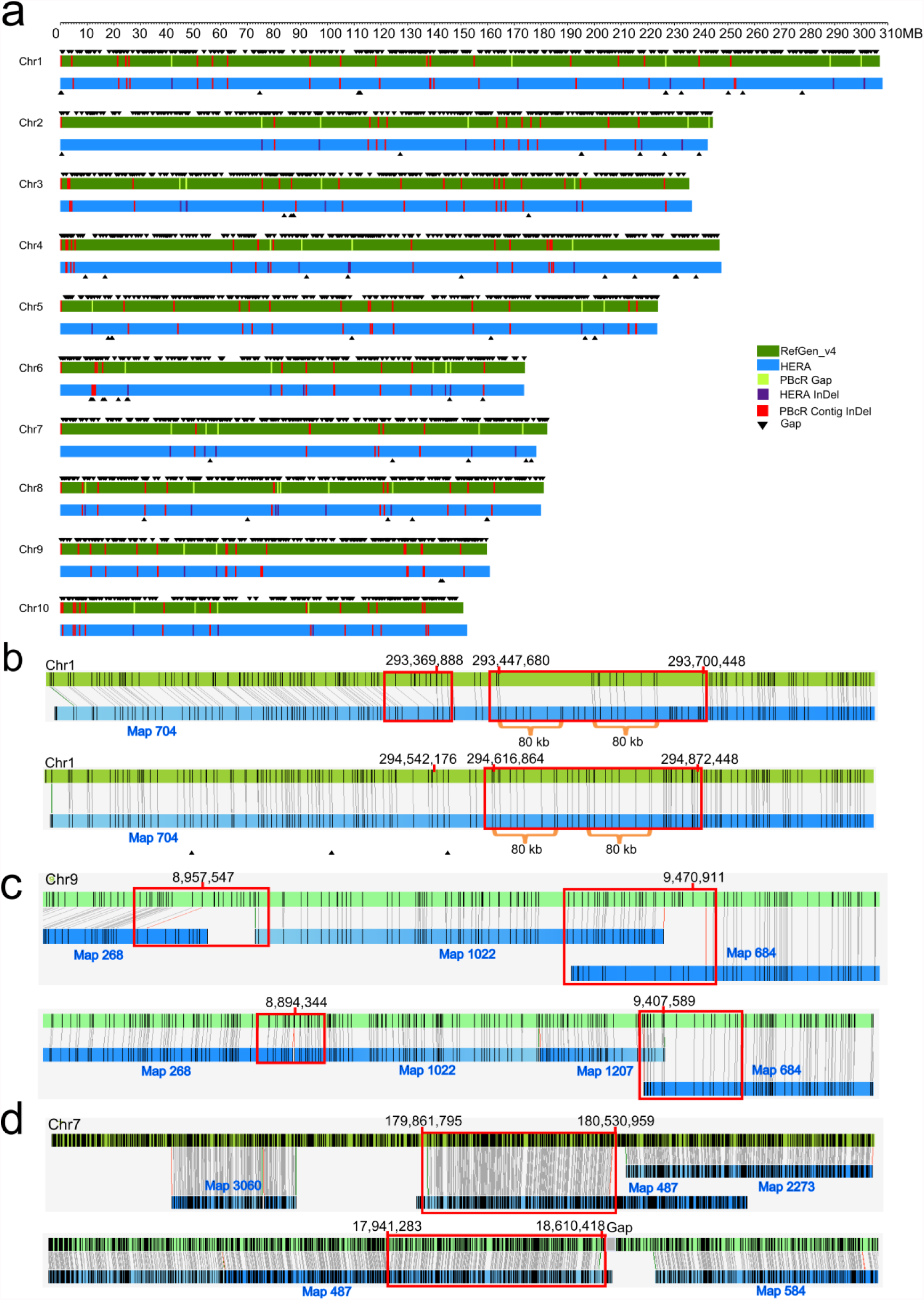
Comparison of maize B73 HERA assembly and B73 RefGen_v4. (**a**) The comparison of HERA assembled B73 genome with the published B73 RefGen_v4^6^. The top green horizontal bar represents RefGen_v4 and the bottom blue horizontal bar represents the HERA assembly. Each black triangle represents a sequence gap. RefGen_v4 contained a total of 2523 gaps while the HERA assembly contained only 76 gaps. The red vertical bars represent the >10 kb indels that were present in the contigs of RefGen_v4. The purple vertical bars represent the >10 kb indels introduced by HERA with the light green vertical bars showing the positions of the corresponding gaps in RefGen_v4. (**b**) An example of HERA assembled sequences that were missed in RefGen_v4. The green horizontal bars represent the maize genome sequences and the blue horizontal bars represent BioNano maps. The upper panel is the alignment between RefGen_v4 and BioNano maps, and the lower panel is the alignment between the HERA assembly and BioNano maps. The gaps (right red box in the upper panel) in RefGen_v4 were filled with ‘N’s. (**c**) An example region with misoriented and missing sequences on maize chromosome 9 in RefGen_v4. Two overhangs were shown in Map 268 and Map 684 in the red boxes, which were caused by a misoriented sequence of 513 kb. In addition, two sequences of 63 kb and 360 kb were missing outside of the misoriented sequence. (**d**) An example of the falsely anchored sequence in RefGen_v4. The BioNano map 487 was partially aligned to two non-adjacent regions in RefGen_v4 while it was aligned to only one region as a whole in the HERA assembly (indicated in the red boxes).

In the HERA assembly, a previously unidentified missing sequence of 387 kb on chromosome 8 close to the centromere in R498_ref1 was added (Supplementary Figure 3). In R498_ref1, there were also 14 known potential indels (inserted or deleted sequences) >10 kb^5^. Using HERA to find paths, we found that at least 8 of the indels could be fixed using a minimum sequence identity of 97% as a cutoff during path extension (Supplementary Figure 3). Almost all of the regions were found to contain tandemly repetitive sequences with multiple peaks in their path length distribution plot. These results also indicated that our assembly method for tandem repeats is valid.

HERA assembled 525 repeat regions with a total size of 6.4 Mb and a max length of 268.7 kb. Among the 495 regions that are covered by BioNano genome maps, 19 (3.83%) of them were found to contain indels >10 kb, setting a max indel error rate of 3.83% for the HERA assembly. As a comparison, there were 20 indels >10 kb also found in the CANU assembled contigs. Although we did not try to fix all of these potential errors, we believe that most or all of them can be fixed under the guidance of BioNano genome maps with known length based on our tests in the tandem repeat regions. These results also demonstrated the power and flexibility of HERA. The quality of the HERA assembled sequences was also validated with both the corrected SMRT reads and Illumina short reads (Supplementary Figure 4). The short reads had a mapping ratio of 97.21%, and covered 99.56% of the HERA assembled regions.

### Improvement of the maize and human HX1 genome assemblies

To demonstrate the power of HERA in assembling more complex genomes, we reassembled the previously published maize B73 genome^6^ and human HX1 genome^4^ that were assembled using SMRT data (Supplementary Table 2). The maize B73 RefGen_v4 assembly contained a large number of small contigs (∼90.55 Mb in total length) that were not anchored to hybrid genome maps and chromosomes, but the hybrid scaffolds contained many unfilled gaps (∼43 Mb in length). After HERA assembly (Supplementary Figure 2), the contig N50 size was increased nearly 50x from 1.3 Mb to 61.2 Mb. The longest contig was 140 Mb, a 20-fold increase from 7 Mb. The total length anchored to chromosomes was increased from 2075.6 Mb to 2104.2 Mb (Supplementary Table 3), and only 2.8 Mb sequences were not anchored. The total number of gaps was reduced from 2,523 to 76 (Figure 3a, 3b). There were 47,838 annotated genes in RefGen_v4^6^ of which 837 were not anchored onto chromosomes. In the HERA assembly, 815 annotated genes were on the contigs that were newly added to chromosomes.

HERA fixed several misarranged sequence errors in RefGen_v4, including two inversions and two translocations (Figure 3c, 3d and Supplementary Figure 5). These errors could not be identified by using BioNano genome maps due to the lack of labelling enzyme recognition sites. These results also indicated that even after intensive curation, the reference genome can still contain structural errors other than indels.

The total number of HERA-assembled regions was 2,632, with a total length of 33.2 Mb and a maximum length of 276.7 kb, including many tandemly repetitive sequences. For example, HERA added a missing tandem repeat of ∼150 kb and another missing sequence of ∼586 kb on chromosome 1 of B73 RefGen_v4 (Supplementary Figure 6). BioNano genome maps covered 2,588 (98.33%) HERA-assembled regions (Supplementary Figure 5f), of which only 47 (1.81%) contained indels >10 kb most likely caused by long tandem repeats (Supplementary Figure 5g). As a comparison, the contigs in RefGen_v4 contained 140 indels >10 kb (Supplementary Table 4), which we did not try to fix. Nonetheless, as shown in the rice genome, we believe that most of these indels can be fixed by HERA.

Similarly, we applied HERA to improve the previously published human genome HX1_FALCON^4^. After HERA assembly (Supplementary Figure 2), the contig N50 size was increased from 8.3 Mb to 54.4 Mb. The longest contig was 109.8 Mb, almost 3 times of the previous max length of 38 Mb. Comparisons between the HERA assembly of HX1, HX1_FALCON and human GRCh38 showed that several gaps in GRCh38 which remained open in HX1_FALCON could be filled in the HERA assembly (Supplementary Figure 7), and that GRCh38 contained some potential errors that could be fixed with the help of HERA (Supplementary Figure 8 and Supplementary Table 5).

### Improvement of the Tartary buckwheat genome assembly

We also applied HERA to assemble the Tartary buckwheat genome. The previously published Tartary buckwheat genome was assembled using only ∼30x SMRT sequence data^17^. In this study, we added more SMRT data to make for a total sequencing depth of ∼70x. We used both PBcR and CANU pipelines to generate two different assemblies, which had a contig N50 size of 0.45 Mb and 1.1 Mb, and a total size of 587.7 Mb and 452.1 Mb, respectively (Supplementary Table 2). After HERA assembly based on the PBcR assembled contigs, the contigs N50 size was increased to 22.24 Mb, with a total genome size of 453.4 Mb, very close to the assembled size of CANU. The hybrid scaffolding with BioNano genome maps resulted in only 12 scaffolds for the entire genome with 5 chromosomes each in one scaffold, two chromosomes each in two scaffolds and one chromosome in 3 scaffolds. Further gap filling with HERA resulted in a contig N50 size of 27.85 Mb and a single-contig chromosome 8. The comparison of the scaffolds with the previously published pseudomolecules^17^ showed that the latter contained a number of misoriented or mispositioned sequences, which obviously resulted from the clustering of short contigs into chromosome sequences by using Hi-C data.

## Discussion

We have reported a highly efficient assembly method, HERA, to resolve repetitive sequences, which is the central objective for all genome assemblers. We tested HERA to improve the genome assembly for rice R498, maize B73 and human HX1 by assembling the previously unassembled repeats including many tandemly repetitive regions. As contig lengths increase, it is more likely for a contig being anchored or positioned correctly onto chromosomes as well as being aligned to BioNano genome maps, and thus the constructed chromosomal sequences contain fewer gaps as well as fewer errors. Highly contiguous and complete genome sequences will increase the accuracy of gene mapping and structural variation identifications when using them as references.

The key difference in dealing with repeats between HERA and SG assemblers such as CANU and FALCON is that HERA assembles a repeat by identifying and scoring the sequence as a whole with different repeat copies being completely separated from each other, while the SG assemblers compress all similar reads above a threshold into one, which makes many different repeat copies unresolvable. In the absence of sequence errors, HERA and SG will generate the same results. However, in the presence of sequence errors which are unavoidable under the current sequencing technologies, the transitive reduction in SG often generates branching nodes from repeat compression, and thus fragments a repeat into many small contigs. HERA is superior in that an overlap graph is reduced to a connection graph in which the repeats are not compressed. HERA is relatively insensitive to unevenly distributed sequencing or self-correction errors. HERA can distinguish two interspersed repeats based on the small difference (<1-2%) at their sequence level by selecting the best paths to connect the true pair of flanking sequences.

In this work, it was not our objective to assemble unique sequences since existing string graph assemblers such as CANU and FALCON accomplish this reasonably well. However, HERA could be applied to assemble unique sequences directly by selecting a non-redundant set of junction reads that are partial unique, partial repeat to replace the assembled contigs in the overlap graph, a strategy that we are testing and has shown very promising results. If a read is aligned to a number of reads without overhangs at both ends and it is also aligned to a number of reads with long overhangs of a few kb at only one end, it can be designated as a junction read. The unique sequences will be assembled by extending on the unique ends. Redundancy of junction reads should be removed to reduce conflicts. The assembly of unique sequences can thus be iterative: the junction reads at different confidence levels based on their mutual sequence similarity are used to generate contigs in multiple steps. Redundancy and chimeric errors in the assembled unique sequences should be further removed before proceeding to the assembly of repetitive sequences. The way an assembler deals with repeats can affect its speed and memory usage, and thus various tradeoffs are often implemented to balance the contig length and program efficiency^1,9,13,18^. By using the method we described here as the second stage in an assembler, the assembly of unique sequences can use a conservative approach by assembling only the high confidence regions in the first stage. We expect that this strategy will decrease the complexity of the program implementation and increase the speed of new assemblers.

Resolution of haplotypes in highly heterozygous genomes is important for studying epigenetic effects on gene expression and functions^19^. It is hard to distinguish heterozygous copies and repetitive sequences when contigs are short. HERA can be used as a phasing tool to resolve haplotypes in the same way as filling the repeat gaps as long as there are no identical sequences longer than reads in the region to be resolved. Furthermore, we noted that there still exist many potential errors besides gaps in the current human reference genome GRCh38. We expect that HERA will be used as a curation tool to help fix many of these errors with SMS data, especially by combining it with BioNano genome maps.

## Methods

### Problems in genome assembly

A DNA sequencer generates random sequence reads of certain lengths. The genome assembly process involves identifying a correct tiling path of overlapping reads and connecting them into contigs (contiguous sequences), eventually recovering the full sequences of each individual chromosome. The difficulty of genome assembly comes from repetitive sequences which cause the tiling paths to be non-linear as shown in SG^8^. There are two major types of repetitive sequences in a genome: tandem repeats and interspersed repeats. Tandem repeats are an array of directly connected highly similar sequence units originating from local sequence duplication. Typical tandem repeats include rDNA and centromeric repeats. Interspersed repeats (including segmental duplications) are non-local similar sequences that are dispersed throughout the genome. In some repeat regions in a genome, both types of repeats can be mixed together to form a long stretch of complex repeats. At present, the SMS reads can reach a N50 length of >15 kb. The repeat regions fully covered by a long read can be easily assembled. What concerns us is the repeats that are longer than read length.

For simplicity, we correct the raw SMS reads to reduce sequence errors, and then we assemble the corrected reads into contigs with existing assemblers. Our objective in this work is to assemble the unassembled regions into contiguous sequences with different repeat copies fully resolved and thus generate ultra-long, up to chromosome-scale, contigs. All the pre-assembled contigs and reads need to be aligned to each other to identify the sequence overlaps. A sequence can be extended by overlapping sequences at both ends if the overlapping sequences are not entirely covered by (contained in) the former. A sequence end can be extended iteratively to form a tiling path of overlapping sequences. Due to sequence errors, the extension of a sequence must allow non-perfect matches. We therefore define a global minimum sequence identity cutoff parameter SI_min_ for filtering out low-confidence overlaps. By allowing the number of overlapping sequences for each sequence to be at most the average sequence depth, the average base accuracy (*α*) of all reads is estimated using the average sequence identity of all high-confidence overlaps. The average error rate in all reads is thus *ε*=1-*α*. SI_min_ is typically set to a value smaller than but close to *α.*

Given a pair of overlapping but non-containing sequences S1 and S2, each contains an overlap region with lengths OL1 and OL2, sequence identity SI, overhang lengths OH1 and OH2, and extension lengths EL1 and EL2. We define the following scores. The overlap score of S1 and S2 is OS = (OL1+OL2)*SI/2. The extension score of S2 extending S1 is ES2 = OS + EL2/2 - (OH1+OH2)/2, and similarly for the extension score of S1 extending S2.

Assume two similar repeat copies R1 and R2 are present between two pairs of unique sequences represented by contig ends C_1_^e^—C_2_^e^ (e is head H or tail T of the contig) and C_3_^e^—C_4_^e^, respectively (Figure 1). Given L_ce_, a sequence end is defined to be sequence that is up to L_ce_ kb from the end. Clearly, due to the sequence similarity of reads for R1 and R2, the contig end C_1_^e^ can be extended to another contig end C_2_^e^, through a tiling path of reads originating from the repeat R1, while it can also be extended to contig end C_4_^e^ by mixing reads originating from both R1 and R2. The objective is to identify the correct path to connect C_1_^e^ to its true neighbor. HERA tries to assemble R1 and R2 separately and generates extended contigs with the process described in the next section.

### Description of the HERA algorithm

#### Overlap graph construction

An *overlap graph, G,* is a colored undirected graph with two node colors and two edge colors. The nodes include anchoring nodes (N^A^) for anchoring sequences, e.g., the pre-assembled contigs, and read nodes (N^R^) for sequencing reads. An E^O^ edge type, which we call an *overlap edge,* represents the overlap between two reads and/or anchoring sequences. An E^C^ edge type, which we call a *coupling edge*, represents a sequence whose ends are represented by two nodes. Each read node is allowed to connect to at most one anchoring node with the highest overlap score. The connection of anchoring nodes can be found by traversing the overlap graph using either depth-first-search or breadth-first-search. A traversal of the graph must obey the following rule: whenever it goes into a node via an overlap edge, it must exit from a coupling edge, and vice versa.

#### A heuristic algorithm for creating paths between anchoring nodes

Starting from a random anchoring node, N^A^_S_, we try to identity all its directly connected anchoring nodes, {N^A^_j_} in *G* and output a set of paths through path traversal/extension. Here we use ‘directly connected’ to indicate that an anchoring node is connected to another one without meeting any other anchoring nodes on the path. Typically, a path extension stops when reaching at a read connecting to another anchoring node, N^A^_j_, i.e., meeting N^A^_j_ on the path. It is computationally expensive to compute all allowed paths emanating from N^A^_S_, and thus we limit the paths chosen for the algorithm. We only consider paths representing sequences whose length is at most a predefined value, denoted as L_ex_. We also require that each read node can be used only once in constructing a path. To maximize the chance for obtaining a valid path between two anchoring nodes, we utilize a combination of fixed scoring schemes and a Monte Carlo approach to construct a set of best-scored paths connecting them.

*Approach I.* In the first extension step, all reads connecting to N^A^_S_ are selected to extend N^A^_S_. For further extension, only the read with the highest overlap score is selected. If scores for two reads are tied, the read with higher sequence identity or the longer read is selected. Note that not all sequences generated in the first step can be eventually extended to another anchoring node. In case of a dead end where no more extending reads can be found, the extension path goes back to the previous node, and then is extended by the next top-scored read.

*Approach II.* Similar to approach I, except that in each extension step, a read with the highest extension score is selected.

*Approach III.* A Monte Carlo approach is used to randomly select a connecting read for each extension. The probability of a connecting read being selected is proportional to its extension score. When the extension stops, a trial for constructing a path is finished, and a new trial can start over. For each starting anchoring node, a limited number of trials may be performed to control the running time.

Finally, a set of paths are created to reach a set of other anchoring nodes {N^A^_j_} from N^A^_S_. Approaches I and II generate a fixed number of paths while the approach III can generate arbitrary number of paths. We require that the approach III generates more paths than approaches I and II. All the paths are put together and duplicated paths are removed to obtain a set of paths from N^A^_S_ to each anchoring nodes N^A^_j_, {P_Sj_}. Obviously, if N^A^_S_ represents a contig end, say C_1_^e^ as before (Figure 1c), and C_2_^e^ is present in the set {N^A^_j_}, then R1 will have a high chance of being present in {P_12_}.

#### Generation of consensus sequences for connecting paths

Among all paths of varied length between a pair of anchoring nodes, we need to find a representative (consensus) sequence as the assembled sequence. If the path lengths are distributed within a small range, such as <10 kb, then all the paths are put into one group. Otherwise, the paths are divided into separate groups based on their length distribution (Figure 2c,d) with each group representing a candidate connecting sequence of different length. Specifically, all sequences were sorted according to their lengths from short to long and divided into groups using a 1 kb window. Comparing each window with the ones immediately before and after it, a window with the largest sum of the path frequencies is designated as a peak window, and if it has the smallest sum of the path frequencies, then it is a valley window. If the lowest path length frequency in the valley window is less than 90% of the highest path length frequency in the right peak window, then the path length of the lowest frequency in the valley windows (left-inclusive) is used to divide the whole set of paths to different groups.

In each group, all paths with length frequency less than half of the highest path length frequency are discarded. The sequences of the remaining paths are aligned to each other and one of the paths with the highest length frequency matching to the largest number of other paths is selected as the consensus sequence. The number of paths matching to the consensus sequence is designated as the valid path number in this group.

When multiple groups of paths are found between a pair of anchoring nodes, a consensus sequence will be generated for each group. The multiple consensus sequences indicate the presence of similar repeat units. If the length of the region is known (e.g. for gap filling), the consensus sequence corresponding to the known length was selected. Otherwise, starting from the group with the highest path length frequency, its valid path number is compared with that of next immediate group with longer paths, iteratively. If there are only two groups or the valid path number in the longer path group is more than half of that in the shorter path group, then the longer path group will be selected. Otherwise, the shorter path group will be selected. Finally the consensus sequence of the selected group is used as the consensus sequence of the region.

If the path lengths in a single group are distributed across a wide window (>100 kb), without a clearly identifiable peak, then the repeat underlying the paths will be viewed as a complex repeat. By default, these kinds of repeats are not used to connect anchoring sequences, i.e., the region is not assembled.

#### Construction of Connection graph and sequence connection

After a consensus sequence is determined between each pair of anchoring nodes, a connection graph (Figure 1d) is constructed on the nodes using the consensus sequence connecting the nodes as edges. The sum of valid path numbers (NP) on both directions between each pair of anchoring nodes is recorded on the edge.

We define a **conflict index** (CI) for each anchoring node. For any anchoring node N^A^_i_, for each of its connecting anchoring nodes {N^A^_j_}, its valid path number is NP_ij_. If two of the connecting anchoring nodes N^A^_m_ and N^A^_n_ have the largest and second largest connecting path numbers NP_im_ and NP_in_, respectively, among all NP_ij_, we define the conflict index for N^A^_i_ as CI_i_^e^=NP_in_/NP_im_. An anchoring node has conflicting connections if its conflict index is larger than a predefined threshold, CI_max_. If an anchoring node N^A^_S_ is connected to other anchoring nodes {N^A^_i_} without conflicting connections, we call the anchoring node in {N^A^_i_} with the largest path number as the matching anchoring node of N^A^_i_.

The anchoring nodes with conflicting connections will not be used in sequence connection unless the conflict is resolved. The non-conflicting anchoring nodes are connected to their matching anchoring nodes with the consensus sequences between them to generate contigs. The consensus sequences without the anchoring sequences from the conflicting connections can be output according to user’s specification.

### SMRT sequencing

The leaf DNA of Tartary buckwheat cultivar Pinku1 was used to prepare SMRTbell libraries (20 kb inserts) with the standard protocol provided by PacBio (Pacific Biosciences, USA), and the sequencing was conducted on a PacBio Sequel system. Sequence reads with a quality score below 0.8 were discarded.

### Assembly of rice, maize, human and Tartary buckwheat genomes

The raw SMRT reads of R498, B73 and HX1 were corrected using the CANU pipeline with default parameters. The corrected reads of R498 were further assembled into sequence contigs using CANU with default parameters. The CANU assembled R498 contigs and the published contigs in B73 RefGen_v4 and HX1 (HX1_FALCON) were used for HERA assembly with the corrected SMRT reads. The raw reads of Pinku1 were assembled using both the CANU and PBcR pipeline with default parameters and the assemblies were improved with HERA. For each genome, the contigs of at least 50 kb were used to construct overlap graphs with the corrected reads. All contigs and the corrected reads were aligned all-against-all with Minimap2 (https://github.com/lh3/minimap2) and BWA^20^ with default parameters to identify the sequence overlaps. To reduce memory usage and computational cost, a reduced overlap graph was constructed. The reads that are fully contained by other reads were discarded and overlaps with sequence similarity <97% were also discarded (SI_min_ set to 97%).

The genomes were assembled using HERA with L_ce_ set to 25 kb, L_ex_ set to 800 kb, and CI_max_ set to 0.75. The HERA assembled super-contigs were combined with BioNano genome maps to generate hybrid maps, and HERA was used again to fill in the gaps in the hybrid maps. The resulting contigs were further connected with HERA. For simplicity, the final order of the HERA assembled super-contigs on chromosomes was determined based on their alignment to reference genome. No further gap filling was performed after the super-contigs were anchored onto chromosomes.

### Cleaning of the Tartary Buckwheat assembly

The HERA assembled Tartary buckwheat contigs were aligned to cpDNA and mtDNA, microbial and human DNAs with BWA-mem and the contigs with alignment length >500 bp were discarded. The remaining contigs between 20-50 kb were aligned to the longer ones and those with sequence coverage >90% were also discarded.

### BioNano map assisted assembly and gap filling

The *de novo* and hybrid assembly of the BioNano genome maps were performed using the IrysView software (BioNano Genomics), using a minimum length of 150 kb as a cutoff. To fill a gap with known length, the path extension was performed with only the Monte Carlo approach, and a path whose length was the closest to the gap length was selected. To fill a gap with unknown length, a standard path extension was performed. The order and group information of the contigs were only used to resolve conflicts.

## Data availability

Rice data (R498) was previously published^5^ with the sequence reads available at GSA database (http://gsa.big.ac.cn/index.jsp) under project PRJCA000313. The maize data was downloaded from NCBI under BioProject number PRJNA10769 and SRA accessions SRX1472849. The human data was downloaded from http://hx1.wglab.org/ and NCBI under BioProject PRJNA301527. The sequence reads of Tartary buckwheat was deposited into GSA database under project PRJCA000402. The HERA assembled genome sequences of B73, HX1, and Pinku1 were deposited into BIG Data Center (http://bigd.big.ac.cn/gwh) under accession numbers GWHAAEN00000000, GWHAAEM00000000 and GWHAAEO00000000. The HERA assembled genome sequences are also available at http://mbkbase.org/B73/, http://mbkbase.org/HX1/ and http://mbkbase.org/Pinku1/.

## Software availability

The HERA software was coded in Perl and is provided as Supplementary Software and is free for academic and research use.

## Acknowledgements

We thank Qiang Gao and Bin Ma in our group, and DeepCamp 8810 team from Lenovo Group Ltd for their help in using the computing clusters to run the software used in this work. We also thank Yan Li in our group for submitting the assembled genome sequences to GenBank. This work was partially supported by the Chinese Academy of Sciences ‘Strategic Priority Research Program’ fund (XDA08020302).

## Contributions

C.L. and H.D. conceived the project and designed the algorithms. H.D. implemented the software and performed the data analysis. C.L. wrote the manuscript with contributions from H.D.

